# Phylogenetic relationships of heteroscleromorph demosponges and the affinity of the genus *Myceliospongia* (Demospongiae *incertae sedis*)

**DOI:** 10.1101/793372

**Authors:** Dennis V. Lavrov, Manuel Maldonado, Thierry Perez, Christine Morrow

**Affiliations:** Department of Ecology, Evolution, and Organismal Biology, Iowa State University; Department of Marine Ecology, Centro de Estudios Avanzados de Blanes (CEAB-CSIC); Institut Méditerranéen de la Biodiversité et d’Ecologie marine et continentale (IMBE), CNRS, Aix-Marseille Université, IRD, Avignon Université; Zoology Department, School of Natural Sciences & Ryan Institute, NUI Galway, University Road, Galway; Ireland and Queen’s University Marine Laboratory, 12–13 The Strand, Portaferry, Northern Ireland

**Keywords:** Porifera, Demospongiae, Heteroscleromorpha, MtDNA, Gene Order

## Abstract

Class Demospongiae – the largest in the phylum Porifera (Sponges) – encompasses 7,581 accepted species across the three recognized subclasses: Keratosa, Verongimorpha, and Heteroscleromorpha. The latter subclass contains the majority of demosponge species and was previously subdivided into subclasses Heteroscleromorpha *sensu stricto* and Haploscleromorpha. The current classification of demosponges is the result of nearly three decades of molecular studies that culminated in a formal proposal of a revised taxonomy (Morrow and Cardenas, 2015). However, because most of the molecular work utilized partial sequences of nuclear rRNA genes, this classification scheme needs to be tested by additional molecular markers. Here we used sequences and gene order data from complete or nearly complete mitochondrial genomes of 117 demosponges (including 60 new sequences determined for this study and 6 assembled from public sources) and three additional partial mt-genomes to test the phylogenetic relationships within demosponges in general and Heteroscleromorpha *sensu stricto* in particular. We also investigated the phylogenetic position of *Myceliospongia araneosa* – a highly unusual demosponge without spicules and spongin fibers, currently classified as Demospongiae *incertae sedis*.

Our results support the sub-class relationship within demosponges and reveal four main clades in Heteroscleromorpha *sensu stricto*: Clade 1 composed of Spongillida, Sphaerocladina, and Scopalinida; Clade 2 composed of Axinellida, Biemnida, Bubarida; Clade 3 composed of Tetractinellida and “Rhizomorina” lithistids; and Clade 4 composed of Agelasida, Polymastida, Clionaida, Suberitida, Poecilosclerida, and Tethyida. The four clades appear to be natural lineages that unite previously defined taxonomic orders. Therefore, if those clades are to be systematically interpreted, they will have the rank of superorders (hence S1-S4). We inferred the following relationships among the newly defined clades: (S1(S2(S3+S4))). Analysis of molecular data from *Myceliospongia araneosa* – first from this species/genus – placed it in S3 as a sister group to *Microscleroderma* sp. and *Leiodermatium* sp. (“Rhizomorina”).

Molecular clock analysis indicated that the origin of the Heteroscleromorpha *sensu stricto* as well as the basal split in this group between S1 and the rest of the superorder go back to Cambrian, while the divergences among the three other superorders occurred in Ordovician (with the 95% standard variation from Late Cambrian to Early Silurian). Furthermore most of the proposed orders within Heteroscleromorpha appear to have middle Paleozoic origin, while crown groups within order date mostly to Paleozoic to Mesozoic transition. We propose that these molecular clock estimates can be used to readjust ranks for some of the higher taxa within Heteroscleromorpha.

In addition to phylogenetic information, we found several unusual mt-genomic features among the sampled species, broadening our understanding of mitochondrial genome evolution in this group and animals in general. In particular, we found mitochondrial introns within *cox2* (first in animals) and *rnl* (first in sponges).

## 1. Introduction

Class Demospongiae Sollas 1895 is by far the largest (>80% of species) and, morphologically, the most diverse in the phylum Porifera (Hooper and Van Soest, 2002). Demosponges are found in both freshwater and marine environments from intertidal zone to abyssal depth and include familiar commercial sponges (Moore, 1908) as well as such oddities as carnivorous sponges (Van Soest et al., 2012). Demosponges fulfill several important roles in benthic ecosystems, being essential players both in the carbon flux (de Goeij et al., 2013) and in the silicon cycle (Maldonado et al., 2012). In addition, sponges have the capacity to modify boundary flow as they pump large volumes of seawater into the water column (Pawlik and McMurray, 2019). With the decline of reef-building corals on tropical reefs, sponges became one of the most important structural elements in these ecosystems and host a variety of other species (Bell, 2008; Bell et al., 2018).

From an evolutionary perspective, sponges — the most likely sister group to the rest of the animals (Simion et al., 2017) — provide insight into the common ancestor of all animals, which, in turn, can help our understanding of the origin of animal multicellularity and evolution of animal body plan (reviewed in Dunn et al., 2015; Renard et al., 2018). Several genomic (Srivastava et al., 2010; Ryu et al., 2016; Borisenko et al., 2016; Francis et al., 2017) and transcriptomic (Riesgo et al., 2014; Guzman and Conaco, 2016) studies of demosponges have been conducted and used to infer steps in animal evolution as well as to clarify various aspects of sponge biology.

*Phylogeny and taxonomy of demosponges*. The relationships among the higher taxa of demosponges have been studied since the second half of the 19th century (reviewed in Cardenas et al., 2012) but are still only partially resolved. Hence, the classification system of the group remains in flux. The most recent pre-molecular taxonomic treatment of the phylum Porifera, *Systema Porifera* (Hooper and van Soest, 2002), subdivided demosponges into three subclasses, and 14 orders, but warned that “resolving the higher systematics of sponges is clearly beyond the capabilities of this present book” (Hooper et al., 2002).

Indeed, the advent of molecular systematics lead to the rejection of many taxa defined based on morphological and embryological data and to the recognition of four major lineages within the class: Keratosa (G1) (Dictyoceratida + Dendroceratida), Verongimorpha (G2) (Chondrosida, Halisarcida, and Verongida), Marine Haplosclerida (G3), and the remaining orders (G4) (at the time, Agelasida, Hadromerida, Halichondrida, Tetractinellida (Astrophorina + Spirophorina), Poecilosclerida, and Spongillina (freshwater Haplosclerida)) (Borchiellini et al., 2004; Lavrov et al., 2008; Sperling et al., 2009; Hill et al., 2013). In addition, a confluence of ultrastructural, embryological and molecular studies lead to the removal of the former order Homosclerophorida from Demospongiae to make a new class Homoscleromorpha.

Recently, a revised classification of the Demospongiae has been proposed that united G3 and G4 into the subclass Heteroscleromorpha and subdivided the latter group into 17 orders (Morrow and Cardenas, 2015). However, this new proposal – now accepted as the framework for the demosponge classification in the World PoriferaDatabase http://www.marinespecies.org/porifera/index.php – was based primarily on nuclear rRNA data and thus needs confirmation from other markers. Furthermore, because many orders within Heteroscleromorpha were defined at the limit of resolving power of molecular markers used in the study (*e.g*., the deepest node with significant support), the relationships among them remained unresolved.

mtDNA sequences are well suited for phylogenetic analysis in Demospongiae as they contain a large amount of data, have a relatively low rate of nucleotide substitutions, and are homogeneous in nucleotide composition (Lavrov and Lang, 2005a; Lavrov et al., 2008; Ma and Yang, 2016; Schuster et al., 2017). Furthermore, in addition to sequence data, mitochondrial gene arrangements can be used to support or refute phylogenetic relationships (Boore and Brown, 1998; Lavrov and Lang, 2005a). Finally, from a technical point of view, the gene-rich nature of animal mtDNA, its stable and nearly identical protein gene content, and the presence of multiple mtDNA copies per cell make mitochondrial genomic research both efficient and cost-effective.

As part of the Porifera Tree of Life project https://portol.org/ we determined mt-genomes from 58 demosponges, including 41 from Heteroscleromorpha *sensu* Cárdenas *et al*. (2012) (aka Heteroscleromorpha *sensu stricto*). Here we report these data and use them to test some of the phylogenetic relationships proposed in the previous studies. Importantly, we included in our dataset *Myceliospongia araneosa* currently classified as Demospongiae *incertae sedis* and resolve its phylogenetic position. In addition, we tested the phylogenetic positions of several demosponge species used for genomic projects and conducted molecular clock analysis of demosponge evolution.

## 2. Materials and Methods

### 2.1. Data collection

#### 2.1.1. Overview

For this project, we PCR amplified, sequenced, assembled, and annotated mt-genomes from 58 demosponges including 41 in the Heteroscleromorpha *sensu stricto*. In addition, we generated low coverage Illumina DNAseq data for 2 species of sponges and assembled mt-genomes from them. Finally, we assembled and annotated mt-genomes from 6 species for which raw genomic and/or transcriptomic data were available. Together with 50 previously published mt-genomes, this resulted in a dataset of 117 complete and nearly complete mt-genomes (Supplementary Table S1). To this dataset we added partial mtDNA sequences from one species of Merliida and one species Desmacellida — the newly proposed orders — as well as a partial mtDNA sequence from an unknown species most closely related to *Plenaster craigi*, resulting in a final dataset of 120 taxa.

#### 2.1.2. Taxon sampling

Species used in this study were chosen to cover much of the suprageneric diversity in Heteroscleromorpha *sensu stricto* (Table 1, Table S1). 17 additional species of demosponges outside of this subclass were sampled and included in the analyses as outgroups (Table 2). Most of the specimens for this study came from the four primary locations: Bocas del Toro in Panama, Irish Sea, Spain’s Mediterranean Sea, and Mo’orea Island in French Polynesia. Additional samples collected elsewhere were provided for this study by April and Malcolm Hill, Alexander Ereskovsky, Gisele Lôbo-Hajdu, Jenna Moore, Michael Nickel, Shirley Pomponi, John Reed, Antonio Solé-Cava, and Gert Wörheide (Tables 1, 2). Finally, 6 mt-genomes were assembled from publicly available DNAseq (*Stylissa carteri*, and *Xestospongia testudinaria*) and RNAseq (*Haliclona (Gellius) amboinensis, Halisarca caerulea, Pleraplysilla spinifera*, and *Scopalina* sp. CDV-2016) data.

**Table 1:**
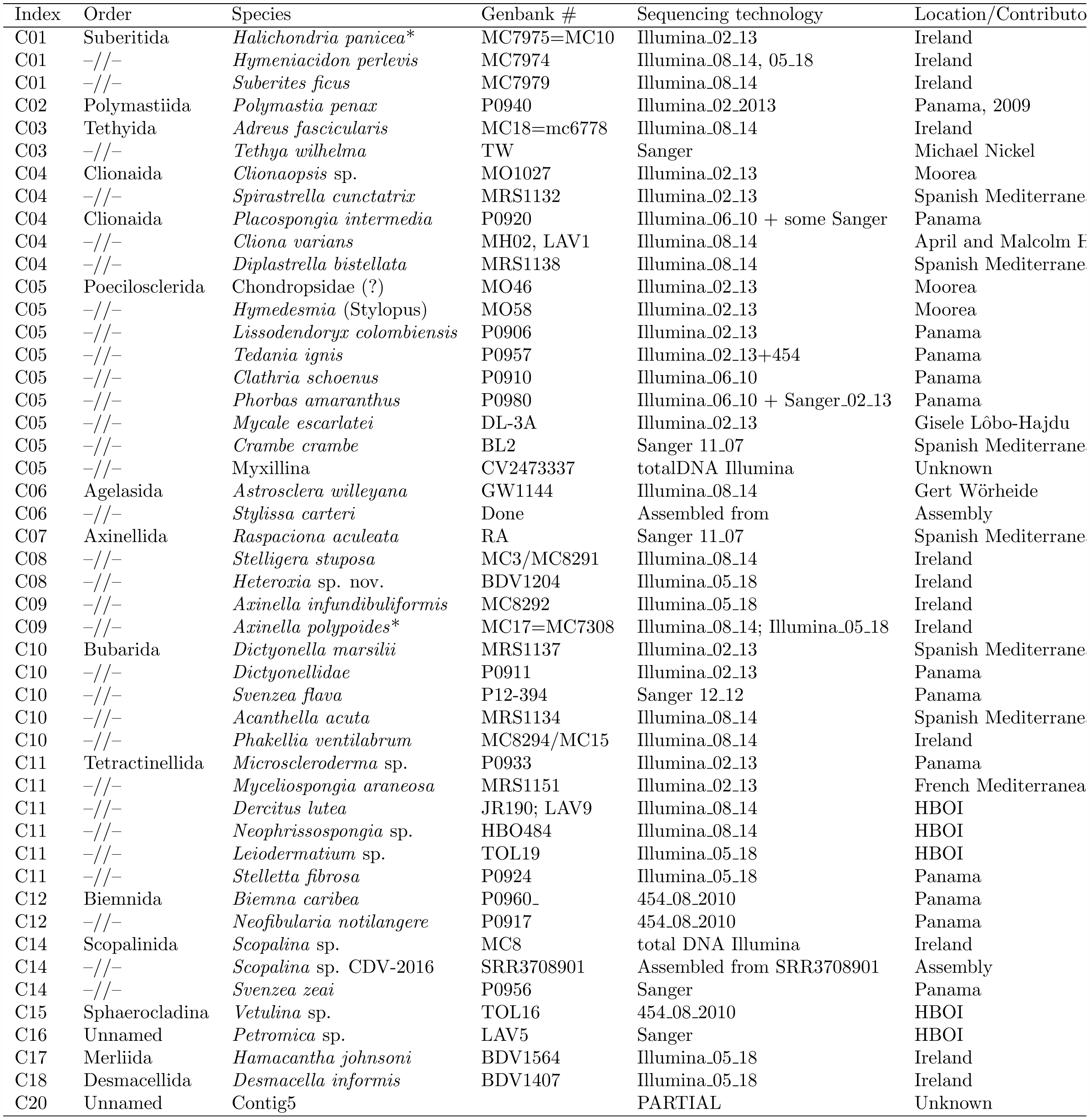
Heteroscleromorph species sampled for this study.

**Table 2:** Other demosponges sampled for the project

| Index | Order | Species | Genbank | Sequencing | Location |
| --- | --- | --- | --- | --- | --- |
| G1 | Dendroceratida | *Darwinella* sp. (simplex) |  | Sanger + Illumina | French Mediterranean |
| G1 | -/- | *Dictyodendrilla dendyi* | DD | Sanger | New Zealand |
| G1 | Dictyoceratida | *Dysidea etheria* FL | DE | Sanger 06,07 | Florida |
| G1 | -/- | *Pleraplysilla spinifera* |  | RNAseq assembly | Assembled from SRR3417588 |
| G1 | -/- | *Carteriospongia foliacens* | GW960 | Illumina_08.14 | Lizard Island, Australia |
| G2 | Chondrosiida | *Chondrosia reniformis* | CR | Sanger | Spanish Mediterranean |
| G2 | Chondrillida | *Thymosia* sp. | MRS1139 | Illumina_2.13_PB | Spanish Mediterranean |
| G2 | -/- | *Halisarca caerulea* | HC | RNAseq assembly | Assembled from SRR5234759 |
| G2 | Verongida | *Aiolochoia crassa* | P0904 | Illumina_02.13 | Panama |
| G2 | -/- | *Hexadella pruvoti* |  | Illumina_08.14 | French Mediterranean |
| G2 | -/- | *Pseudoceratinidae* sp. | MO1014 | Illumina_08.14 | Moorea |
| G3 | Haplosclerina | *Chalinula loosanoffi* | CLOOS | Illumina_08.14 | Virginia, USA |
| G3 | -/- | *Haliclona implexiformis* | SI06-32 | Sanger-07 | Panama |
| G3 | -/- | *Haliclona indistincta* | MC7978 | Sanger-12 | Ireland |
| G3 | -/- | *Haliclona manglaris* | P0905 | Illumina_HiSeq_08.10 | Panama |
| G3 | -/- | *Haliclona verrucosa* (?) | MRS1163 | Illumina_02.13 | French Mediterranean |
| G3 | -/- | *Niphates erecta* | P0902 | Illumina_06.10 | Panama |
| G3 | -/- | *Niphates erecta* | FL06507 | Sanger-12 | Florida |
| G3 | Petrosina | *Neopetrosia proxima* | P0945 | Illumina_02.13 | Panama |
| G3 | -/- | *Xestospongia testudiaris* | XT | Contigs assembly | http://xt.reefgenomics.org |

#### 2.1.3. DNA extraction, PCR amplification

Collected sponge samples were preserved in either 95% ethanol or 3M GuCl solution. Total DNA was extracted with a phenol-chloroform method modified from (Saghai-Maroof et al., 1984). Porifera-optimized conserved primers developed in our laboratory (Lavrov et al., 2008) were used to amplify short (400–1000 nucleotide) fragments of several mitochondrial genes for each species. Two species-specific primers were designed for each of these genes for PCR amplification. Complete mtDNA was amplified in several overlapping fragments using the Long and Accurate (LA) PCR kit from TAKARA.

#### 2.1.4. Sequencing

Three sequencing technologies were utilized in the project (Table 1, Table 2). MtDNA sequences from 11 species were determined using Sanger method (Sanger et al., 1977). For each of these species all LA-PCR fragments were combined in equimolar concentrations, sheared into pieces 1–2 kb in size and cloned using the TOPO Shotgun Subcloning Kit from Invitrogen. Colonies containing inserts were collected, grown overnight in 96-well blocks and submitted to the DNA Sequencing and Synthesis Facility of the ISU Office of Biotechnology for high-throughput plasmid preparation and sequencing. Gaps in the assembly were filled by primer-walking.

MtDNA sequences from three species were determined using 454’s sequencing technology. PCR reactions for each species were combined in equimolar concentration, sheared and barcoded as described in Gazave et al. (Gazave et al., 2010). Barcoded PCR fragments were combined together and used for the GS FLX Titanium library preparation (454 Life Sciences). Pyrosequencing was carried out on a Genome Sequencer FLX Instrument (454 Life Sciences) at the University of Indiana Center for Genomics and Bioinformatics.

Finally, mtDNA sequences for 46 species were determined by using the Illumina technology. For these species, PCR reactions for each species were combined in equimolar concentration, sheared and combined together with or without barcoding. Libraries were prepared using the Illumina TruSeq DNA PCR-Free Library Prep Kits. Sequencing was carried on MySeq and HiSeq instruments at the DNA Sequencing and Synthesis Facility of the ISU Office of Biotechnology.

#### 2.1.5. Sequence assembly

Different assemblers were used depending on the type of data collected. The STADEN package v. 1.6.0 (Staden, 1996) with Phred basecaller (Ewing and Green, 1998; Ewing et al., 1998) was used to assemble the Sanger sequences. Abyss (Simpson et al., 2009), Mira (Chevreux et al., 1999), PCAP (Huang et al., 2003), and SPAdes (Bankevich et al., 2012) were used to assemble 454 and Illumina sequences (Table S1). In most cases, several programs were used for the assembly and results compared and compiled together. When barcodes were used, sequences were first separated by the barcode. When barcodes were not used, species selection was carried out to exclude closely related species from the same library. PCR and Sanger sequencing was used to resolve any ambiguities.

#### 2.1.6. Sequence annotation

We used flip v. 2.1.1 (http://megasun.bch.umontreal.ca/ogmp/ogmpid.html) to predict ORFs in assembled sequences; similarity searches in local databases and in GenBank using FASTA (Pearson, 1994) and NCBI BLAST network service (Benson et al., 2008), respectively, to identify them. Proteincoding genes were aligned with their homologues from other species and their 5’ and 3’ ends inspected for alternative start and stop codons. Genes for small and large subunit ribosomal RNAs (*rns* and *rnl*, respectively) were identified based on their similarity to homologous genes in other species, and their 5’ and 3’ ends were predicted based on sequence and secondary structure conservation. Transfer RNA genes were identified by the tRNAscan-SE program (Lowe and Eddy, 1997). RNAweasel (Lang et al., 2007) was used to search for introns in coding sequences. The exact positions of introns were adjusted based on alignments of coding sequences that contained them.

### 2.2. Phylogenetic inference

#### 2.2.1. Phylogenetic analysis based on mitochondrial coding sequences

Inferred amino acid sequences of individual mitochondrial proteins were aligned with Mafft v6.861b (Katoh and Standley, 2013). Conserved blocks within the alignments were selected with Gblocks 0.91b (Talavera and Castresana, 2007) using relaxed parameters (parameters 1 and 2 = 0.5, parameter 3 = 8, parameter 4=5, all gap positions in parameter 5). Cleaned alignments were concatenated into a supermatrix containing 3,626 amino acid positions for 120 species. This alignment was analyzed with PhyloBayes MPI 1.7 (Lartillot et al., 2013) under the CAT+G4 model, until maxdiff = 0.264969, meandiff = 0.00650579 (∼ 12,000 cycles). The chains were sampled every 10th tree after the first 1000 burn-in cycles to calculate the consensus tree.

#### 2.2.2. Gene order analysis

Mitochondrial gene orders were converted to gene adjacency matrices using the *gogo* program (https://github.com/dlavrov/bio-geneorder, unpublished). The matrices were further modified as required by TNT and RAxML and used in these programs to infer the Maximum Parsimony (MP) and Maximum Likelihood (ML) trees, accordingly. For the parsimony analysis, we tried both the traditional (*i.e*., random addition of sequences + TBR branch swapping followed by additional branch swapping of trees in memory) and “new technology” (*i.e*., with with ratchet, tree-drifting, and tree-fusing followed by additional branch swapping) strategies implemented in TNT (Goloboff et al., 2008). Because the same set of trees was retained in both analyses, we conducted bootstrap analysis using only the first approach. For the ML analysis, we used the multistate model in RAxML-NG (Kozlov et al., 2018): “raxml-ng --all --msa 68taxa.phy --model MULTI13 MK.” To check for the effect of more frequent tRNA rearrangements in animal mitochondrial genomes, we created an alternative adjacency matrix based, where position of each gene was recorded relative to the position of the closest major (*i.e*., protein or rRNA) upstream and downstream genes and repeated the MP and ML analyses.

### 2.3. Molecular clock analysis

PhyloBayes 4.1c (Lartillot et al., 2009) was used for the molecular clock analysis on the fixed tree topology inferred from mitochondrial coding sequences (Fig. 2). Two chains were run for 40,000 cycles and convergence was assessed by estimating discrepancies and effective sizes for continuous variables in the model based on the “acceptable run” criteria: maxdiff <0.3 and minimum effective size >50. We used the default log-normal autocorrelated relaxed clock model (-ln), a generalized time-reversible (GTR) amino acid substitution matrix (-gtr), a Dirichlet mixture profile (-cat), and a discrete gamma distribution with four categories-dgam [4].

**Fig. 1.**
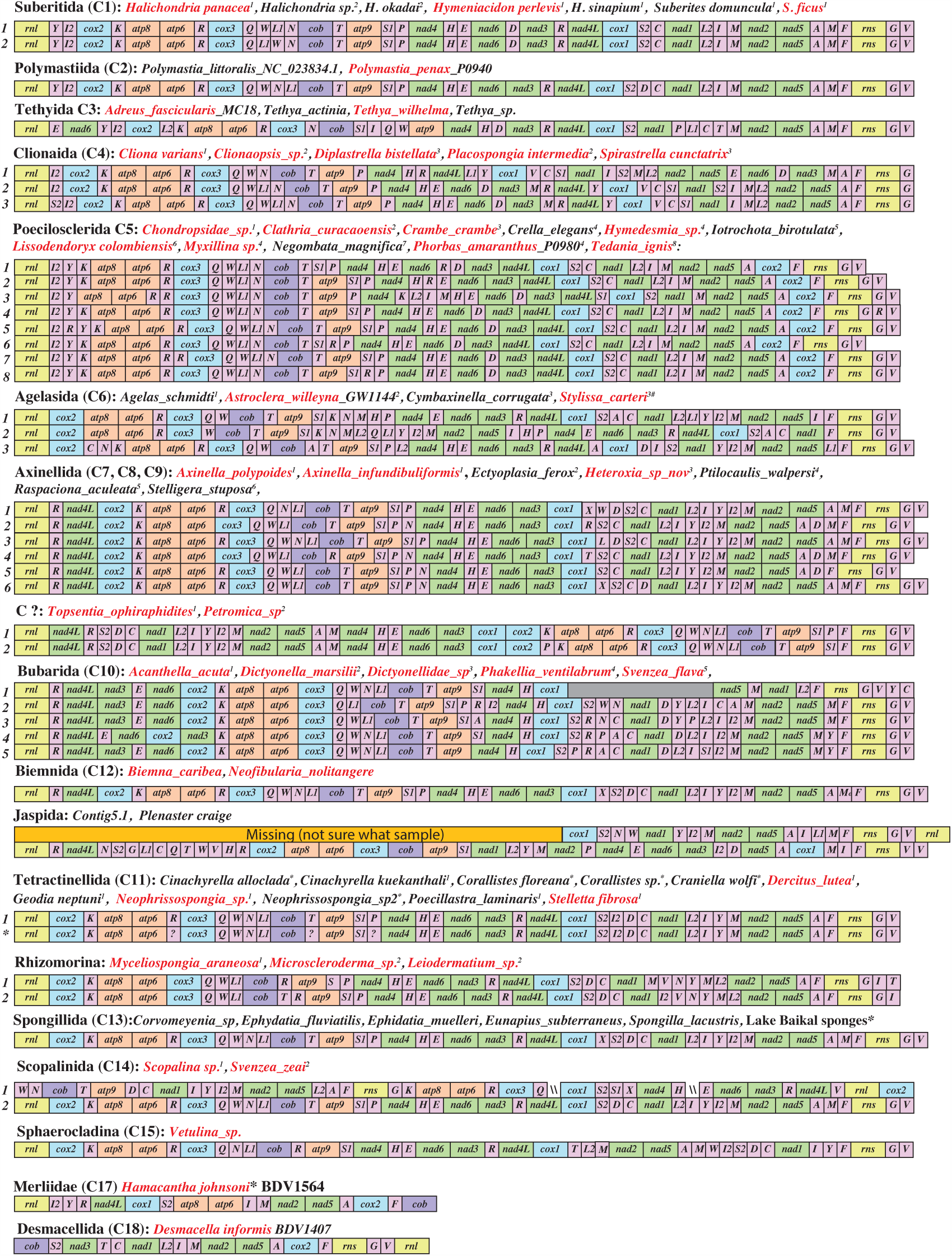
Mitochondrial gene arrangements in Heteroscleromorpha *sensu stricto*. Protein and rRNA genes (larger boxes) are: *atp6*, *8-9* – subunits 6, 8 and 9 of the F0 ATPase, *cox1-3* – cytochrome *c* oxidase subunits 1-3, *cob* – apocytochrome *b* (*cob*), *nad1-6* and *nad4L* – NADH dehydrogenase subunits 1-6 and 4L, *rns* and *rnl* – small and large subunit rRNAs. tRNA genes (smaller boxes) are abbreviated using the one-letter amino acid code. The two arginine, isoleucine, leucine, and serine tRNA genes are differentiated by numbers with *trnR(ucg)* marked as *R1*, *trnR(ucu)* – as *R2*, *trnI(gau)* – as *I1*, *trnI(cau)* – as *I2*, *trnL(uag)* – as *L1*, *trnL(uaa)* as *L2*, *trnS(ucu)* – as *S1*, and *trnS(uga)* – as *S2*. All genes are transcribed from left to right. Genes are not drawn to scale and intergenic regions are not shown.

**Fig. 2.**
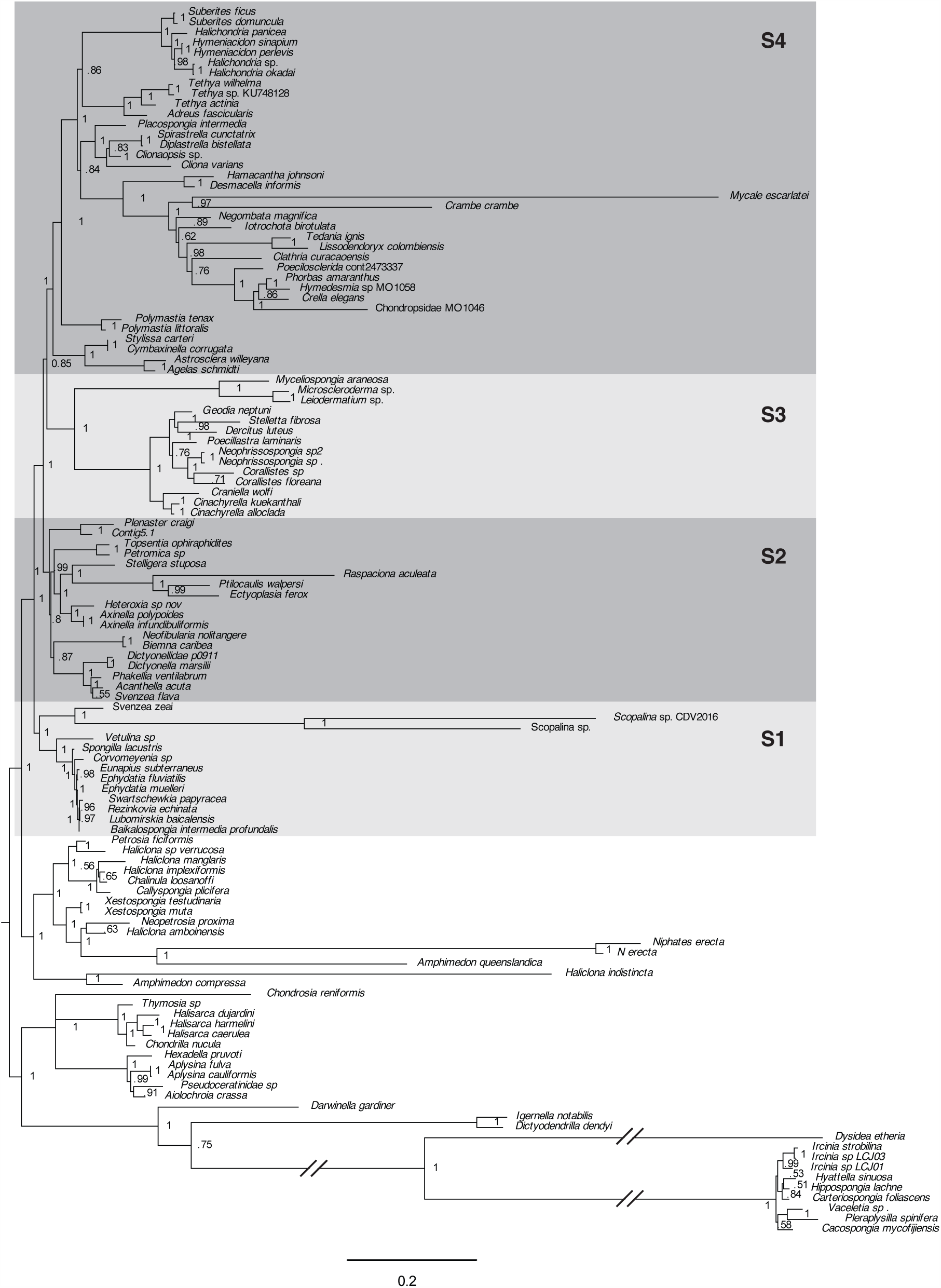
Posterior majority-rule consensus tree obtained from the analysis of concatenated mitochondrial amino acid sequences (3,626 positions) under the CAT+GTR+Γ model in the PhyloBayes-MPI program. The number at each node represents the Bayesian posterior probability. The branches marked by a broken line symbols are shown half of their actual lenghts.

We used three calibration points for the analysis. We put a uniform prior on the root of the demosponges (split between G1+G2 and G3+G4) between 541 and 515 MA (beginning of Cambrian – crown-group heteroscleromorph fossil (Botting et al., 2015)). We constrained the split between freshwater sponges and *Vetulina* sp. at between 410 and 298 MY (the lower bound is defined by the observation that species diversification in lakes prior to the Devonian was limited by low nutrient loads and high sediment loads (Cohen, 2003), the upper bound is defined by the first reported freshwater sponge fossil (Schindler et al., 2008)). Finally, we constrained the origin of the crown group Baikal sponges (the split between *Baikalospongia intermedia profundalis* and *Lubomirskia baicalensis*) between 30 and 6 MY based on Lake Baikal history and fossil record of Lake Baikal sponges (Veynberg, 2009). Although several other fossils have been used in a recent study (Schuster et al., 2018b), the ones used here appear to be the least controversial (but see discussion). All calibration ranges were specified as soft bounds (-sb option), which allocates 0.025 of the total probability outside the specified bounds. Dates were assessed by running readdiv with 250 generations removed as burn-in for each analysis.

## 3. Results

### 3.1. Genome organization and evolution

#### 3.1.1. Genome organization and gene content

Most analyzed mtDNAs of demosponges were circular-mapping molecules, each containing a conserved set of 14 protein-coding, two ribosomal RNA (rRNA) and 24 or 25 tRNA genes. However, the following exceptions to this typical organization have been found in the newly sequenced mt-genomes:

1. the mitochondrial genome of the poecilosclerid *Mycale escarlatei* did not assemble into a single circular molecule. Instead, several alternative arrangements have been found for most mitochondrial genes, indicating an unusual and likely multi-chromosomal genome architecture. Preliminary data from several other species in the genus *Mycale* suggested a similar organization (unpublished).
2. the mitochondrial genome of the *Scopalina* sp. assembled into three contigs with AT-rich sequences at the ends of each of them. The contig containing *cox1* had twice the coverage than the other two. It is not clear whether these contigs represent individual chromosomes or are results of a genome duplication/mis-assembly.
3. *atp9* was missing in two poecilosclerids: *Lissodendoryx* sp. and an chondropsidae species MO1046 as well as haplosclerids *Amphimedon queenslandica* (Erpenbeck et al., 2007) and *Neopetrosia proxima*. Neither *atp9* nor *atp8* was found in mtDNA of the two sampled representatives of *Niphates erecta*.
4. Some variation in tRNA gene content was observed within Heteroscleromorpha, including the loss of *trnC(gca)* in *Crambe crambe*, the loss of *trnD(guc)* in *Astroclera willeyana* and closely related *Agelas schmidtii*, as well as *Clathria curacaoensis*, potential loss of *trnQ(uug)* in two *Corallistes* species, and a potential loss of two mt-tRNA genes in *Scopalina* spp. More variation in tRNA gene content was found in other demosponge subclasses. In addition to the previously reported loss of all but two tRNAs in Keratosa, we observed losses of multiple tRNA genes in *Chondrosia reniformis* Chondrosiidae, Verongimorpha) and several representatives of the clade B of haplosclerids.
5. A few unusual and/or redundant tRNAs have also been observed in newly sequenced mt-genomes:
  a. *trnI(aau)* instead of the usual *trnI(gau)* was found in *Adreus fascicularis*;
  b. *trnR(acg)* instead of *trnR(ucg)* was found in *Cliona varians*;
  c. *trnL(caa)*, a third gene for Leucine tRNA was found in *Heteroxia beauforti*;
  d. *trnY(aua)*, in addition to *trnY(gua)*, was found in *Negombata magnifica*
  e. unusual *trnX(uua)* that would be predicted to read the stop codon UAA was found in *Stelligera stuposa* but had the lowest cove score among all tRNAs in this species;
  f. *trnA(ugc)* was present twice in *Stylissa carteri* as in closely related *Cymbaxinella corrugata*, where it was shown to be recruited from *trnT(ugu)* (Lavrov and Lang, 2005b);
  g. *trnR(ucu)* was present twice in *Vetulina* sp.

Most (78 out of 117) sampled complete demosponge mitochondrial genomes were in 16-22 kbp size range, which a mean size of ∼20.5 kbp. However, a few mitochondrial genomes were larger in size (>30kpb in *Dysidea etheria*), mostly due to the expansions of non-coding regions. All analyzed mitochondrial genomes had similar nucleotide composition (A+T content between 56-74%) and, with two exceptions, displayed overall negative AT- and positive GC-skews of the coding strand (the two exception were *Dysidea etheria* with AT-skew=0.01 and *Scopalina* sp.1, with AT-skew=0.03).

#### 3.1.2. Gene order

Mt-genomes in Heteroscleromorpha *sensu stricta* shared with each other at least six gene boundaries, with the mean number of shared boundaries between a particular species and the rest of heteroscleromorphs varying between 10 for *Plenaster craigi* and nearly 29 for *Svenzea zeai* and freshwater sponges. Most of the differences in mitochondrial gene orders were caused by transpositions of tRNA genes. However, rearrangements of “major” (protein and rRNA) genes were also present. Interestingly, no inversions were found in Heteroscleromorpha mtDNA: all genes had the same transcriptional polarity. Our previous study (Lavrov and Lang, 2005a) demonstrated that two randomly reshuffled mt-genomes with all genes transcribed from the same strand would share on average only one gene boundary and that the probability for sharing more than three boundaries by chance is less than 0.05. Thus, mtgene arrangements in Heteroscleromorpha contain a detactable phylogenetic signal, which we utilized in the phylogenetic analysis (see below).

#### 3.1.3. Introns in cox1, cox2, and rnl

In total, 16 *cox1* introns have been found in nine heteroscleromorphs sampled for this study: one in *Adreus fascicularis* (order Tethyida), two in each *Axinella polypoides* and *Axinella infundibuliformis* (order Axinellida), two in *Cliona varians* (order Clionaida), one in *Phakellia ventilabrum* (order Bubarida), two in *“Svenzea” flava* (classified as Scopalinida, but – based on our data – closely related to *Acanthella acuta* and *P. ventilabrum* (order Bubarida)), three in *Leiodermatium* sp., one in *Microscleroderma sp*., and two in *Myceliospongia araneosa*. In addition, two *cox1* introns were found in the Verongimorpha *Thymosia* sp., doubling the number of previously known demosponge orders with mt-introns. In three species (*C. varians, Leiodermatium* sp. and *S. flava*) group I intron was found in position 387, previously reported only in the tetractinellid *Stupenda singularis* (Kelly and Cárdenas, 2016). In *Myceliospongia araneosa* an intron was found in position 714, as previously reported for some *Cinachyrella* species (Schuster et al., 2017). An intron was found in position 723 in four species: *A. polypoides, C. varians, Microscleroderma* sp., and *Leiodermatium* sp. An intron was found in position 870 in five species: *Adreus fascicularis, Axinella polypoides, Thymosia* sp., *Leiodermatium* sp. Finally, in three species an intron was found in position 1141 (*P. ventilabrum, S. flava*, and *Thymosia* sp.).

Unexpectedly, we also found group II (domain V) introns in two other mitochondrial genes: *cox2* of *Acanthella acuta* and *rnl* of *Dictyonella marsilii* and another Dictyonellidae species (P0911). The *cox2* intron in *Acanthella acuta* contained an ORF most similar to the group II intron reverse transcriptase/maturase of a microalga *Ulva ohnoi*. It should be noted, that *A. acuta* likely contains another intron in *cox1*, but the sequence of this region remains undetermined. The *rnl* intron in the two Dictyonellidae species was found in the same position as in the three placozoan species (Burger et al., 2009) and contained a large region that displayed high sequence similarity to a region within mt-lrRNA gene in those species.

#### 3.1.4. Rate of sequence evolution

Most of the demosponge mitochondrial genomes displayed a relatively low rate of sequence evolution. However, significant acceleration in the rates of evolution occurred in several lineages of demosponges, most noticeably Dendroceratida, but also clade B of Haplosclerida, genus *Scopalina*, and genus *Mycale* (Fig. 2). Interestingly, in the latter two groups acceleration in the rates of sequence evolution is associated with the likely unusual mt-genome architecture.

### 3.2. Phylogenetic analysis based on a supermatrix of inferred amino acid sequences

Bayesian phylogenetic analysis based on concatenated amino-acid sequences of mitochondrial protein genes from 120 species of demosponges produced a tree of demosponge relationships consistent with the four major groups suggested by Borchiellini et al. 2004: Keratosa (G1), Verongimorpha (=Myxospongiae) (G2), marine Haplosclerida (G3) and Heteroscleromorpha *sensu stricto* (G4). Although, no outgroups were included in our analysis, we rooted the tree between the two former and two later clades, *e.g*., ((G1 + G2)(G3 + G4)) according to the results of the previous studies (Borchiellini et al., 2004; Lavrov et al., 2008; Simion et al., 2017). Within Heterosleromorpha, most orders proposed by Morrow and Cardenas 2015 were recovered as monophyletic groups, although, based on a limited species sampling. A few cases in which a particular species did not place within their accepted orders (*e.g*., *Topsentia ophiraphidites* and *Svenzea flava*) were likely due to their erroneous classification (see Discussion).

Within Heteroscleromorpha *sensu stricto*, we found support for four larger clades or superorders (named here S1–S4). First, marine orders Scopalinida and Sphaerocladina (represented by *Vetulina* sp.) grouped with freshwater sponges (order Spongillida) and together they formed the sister group to the rest of the taxa.

Second, three large lineages were supported among the remaining heteroscleromorphs. The first of them (S2) contained Axinellida, Biemnida, and Bubarida, along with *Topsentia ophiraphidites* and *Petromica* sp. The second group (S3) comprised Tetractinellida: Astrophorina, Spirophorina, Rhizomorina + *Myceliospongia araneosa*. The last group (S4) included Agelasida, Clionaida, Poecilosclerida, Polymastida, Suberitida, and Tethyida. The latter two groups were sister lineages in the phylogenetic tree. While the support for each of these lineages was high, their interrelationship was only moderately supported.

Three additional small orders (Merliida, Desmacellida, and Trachycladida) have been proposed within the Heteroscleromorpha (Morrow and Cardenas, 2015). We were able to obtain partial sequences from representatives of two of these orders: *Hamacantha johnsoni* (Merliida) and *Desmacella informis* (Desmacellida). Both of these species grouped together in our phylogenetic tree and formed a sister group to Poecilosclerida.

#### 3.2.1. Phylogenetic position of emerging sponge genomic model systems

Our dataset included nine species of sponges for which high throughput DNA and/or RNA data were available and which, therefore, could be considered as emerging sponge model systems. Although the phylogenetic position of most of these species was as expected, *“Stylissa carteri”* grouped closely with *Cymbaxinella corrugata* with the coding sequences being >99% identical between the two species, indicating a likely misidentification of the sponge. Furthermore, *Plenaster craigi*, grouped with the sponges from the orders Axinellida, Biemnida, and Bubarida in sequence-based phylogenetic reconstructions, but was not placed in any of these groups. Thus, it should not be consider a representative of Axinellida (Lim et al., 2017) and its phylogenetic position should be further investigated. Finally, we note that there was a significant level of cross-contamination between by *Xestospongia testudinaria* DNA of *Stylissa carteri* such that both mitochondrial genomes can be assembled from either species sequencing library.

### 3.3. Phylogenetic analysis based on gene order data

To further investigate phylogenetic relationships within Heteroscleromorpha, we extracted gene order data from complete or nearly complete mtgenomes from this group and used them for ML analysis in RAxML-NG (Kozlov et al., 2018) and parsimony analysis in TNT (Goloboff et al., 2008). The results of these analyses (Fig. 3, Fig. S1) generally support the heteroscleromorph relationships reconstructed from sequence data, including its subdivision into major superorders and the phylogenetic position of *Myceliospongia araneosa*. However, three major discrepancies were found:

**Fig. 3.**
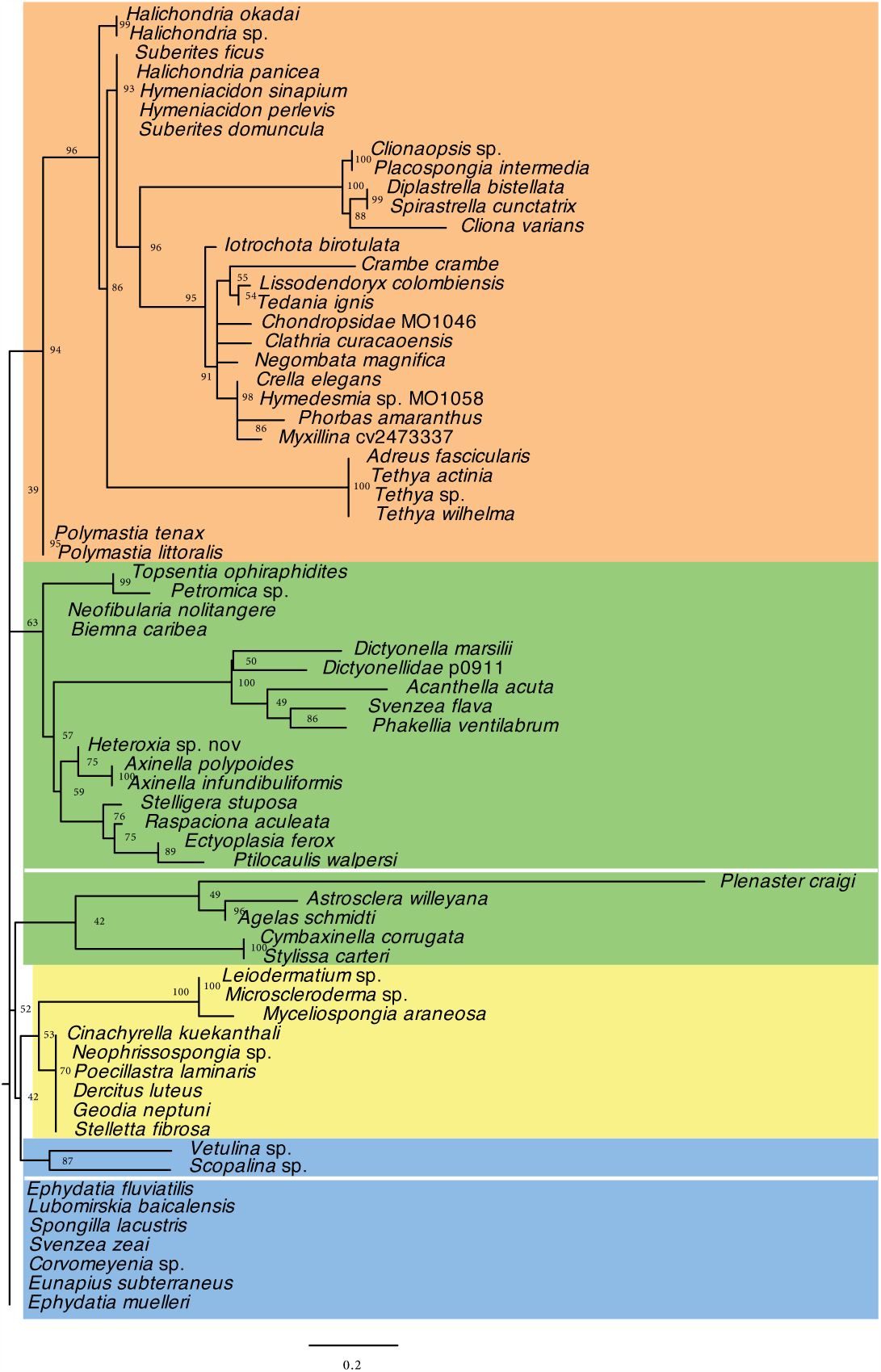
Maximum Likelihood tree inferred from the analysis of mitochondrial gene order data in Heteroscleromorpha. Gene boundaries were encoded as multistate characters and analyzed in RAxML-NG under the MULTI13 MK model. Numbers above branches show the bootstrap support.

First, S1 superorder (freshwater sponges, *Vetulina* sp. and Scopalinida) was not reconstructed as a monophyletic group. Instead *Vetulina* sp. and *Scopalina* sp. either grouped with Tetractinelida plus *Myceliospongia araneosa* or was a part of a polytomy at the base of the tree. The lack of support for S1 superorder should not be surprising, given that the gene order in freshwater sponges was inferred to be ancestral for the Heteroscleromorpha (Lavrov et al., 2008).

Second, phylogenetic position of the order Agelasida was unstable, as it either grouped with S3 + *Vetulina* sp. and *Scopalina* sp. (ML analysis) or its position was unresolved among the superorders.

Finally, *Plenaster craigi* grouped with *Agelas schmidti* and *Astrosclera willeyana* within Agelasida. However, as noted above, *Plenaster craigi* has the most derived mitochondrial gene order, reflected in its by far the longest branch in the ML tree.

To check if any inconsistencies between the sequenced-based and gene order-based phylogenies were due to saturation of phylogenetic signal in gene order data caused by frequent movement of tRNA genes, we repeated both ML and MP analyses using the mulstistate encoding based on closest major (protein or rRNA) genes (see Lavrov and Lang, 2005a, for details). Indeed, *Plenaster craigi* grouped with S2 taxa in both of these analyses. However, the position of Agelasida was still unresolved and there was no support for the monophyly of Tetractinellida, although *Myceliospongia araneosa* still grouped with *Microscleroderma* sp., and *Leiodermatium* sp.

Finally, we visually identified several unique rearrangements associated with some large lineages of Heteroscleromorpha. In particular, we found that the sampled representatives of the orders Axinellida, Bubarida, Biemnida, along with *Topsentia ophiraphidites* and *Petromica* sp. shared a translocation of *trnR-nad4L* downstream of *rnl*, while orders Suberitida, Polymastiida, Poecilosclerida, and Clionaida shared a translocation of *trnY* and *trnI2* to the gene boundary downstream of *rnl*. In addition, all Poecilosclerida species showed a translocation of *cox2* into the gene junction between *nad5* and *rns*, while all Clionaida species had *trnV* moved immediately downstream of *cox1* from its conserved position between *trnG* and *rnl*.

### 3.4. Molecular clock analysis

We used the same dataset as for the sequence-based phylogenetic analysis as well as the consensus tree inferred in that analysis to estimate the time of divergences among major lineages of demosponges (Fig. 4, Fig. S2). Focusing on heteroscleromorph sponges, we estimated that the split between Haploscleromorpha and Heteroscleromorpha *sensu stricto* occurred in Cambrian between 536 and 481 MYA, the basal split within Heteroscleromorpha in Cambrian or Early Ordovician between 506 and 445 MY, the splits among three other superorders occurred in Ordovician, while most of the orders of demosponges have Ordovician to Devonian origins. By contrast, basal splits within most orders (that can be used as a proxy for the crown group) occurred from Early Permian to Late Triassic. Several exceptions to this pattern have been found. First, we noted that the split between *Myceliospongia araneosa, Microscleroderma* sp., and *Leiodermatium* sp. with the rest of Tetractinellida occurred between 456 and 385 MYA, corresponding to the time of the origin of most orders. Second, the basal split within Axinellida (e.g., between Raspaillidae + Stelligeridae and Axinellidae) is estimated to occur at similar time, between 444 and 361 MYA. By contrast, *Hamacantha* (order Merliida) and *Desmacella* (order Desmacellida) estimated to split only between 277 and 102 MYA and to split from Poecilosclerida 375 and 287 MYA. We suggest that these discrepancies can be used to re-define several orders in Heteroscleromorpha *sensu stricto* (see Discussion).

**Fig. 4.**
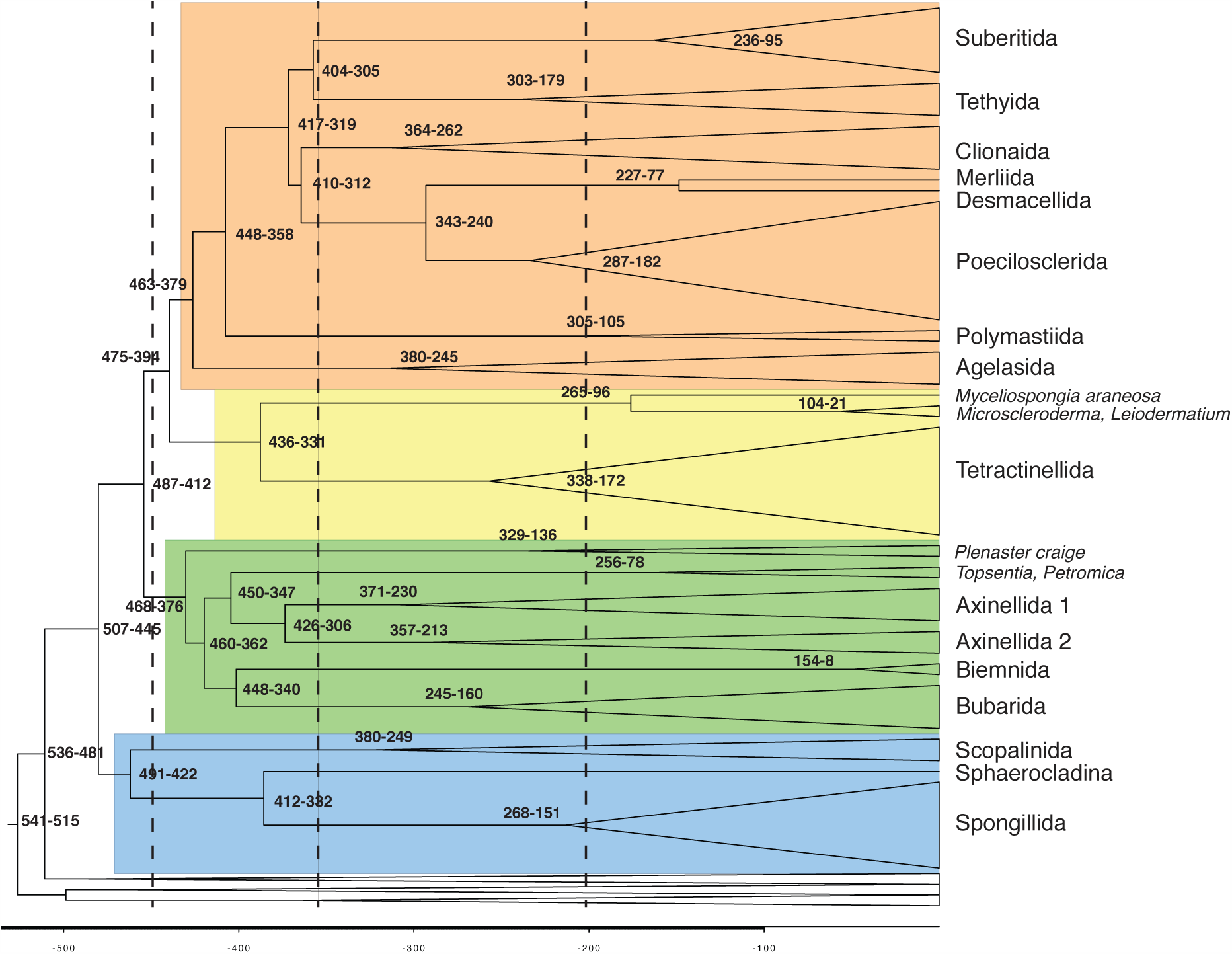
Simplified time calibrated phylogeny of Demospongiae based on Bayesian analysis in PhyloBayes. Only heterosleromorph orders are labeled. Numbers at internal nodes indicate their upper and lower age limits. The four superorders within Heteroscleromorpha are shaded in colors. The full tree is shown in Figure S1.

## 4. Discussion

### 4.1. Assembling the new dataset of mtDNA data for demosponges

Despite rapid progress in sequencing technologies, sequence data remain scarce for many taxa of marine animals, including demosponges. When such data are available, they are often limited to partial sequences of individual genes, commonly those for nuclear rRNA and/or mitochondrial *cox1*. While even these partial data have been instrumental for our re-evaluation of demo-sponge relationships, larger datasets are needed to test proposed phylogenetic hypotheses. Ultimately, a representative sampling of genomic loci should be a dataset of choice in any phylogenetic analysis. However, such datasets are available for only a few species of sponges. Furthermore, assembling and analyzing genomic dataset present various challenges, especially in sponges, where contamination by foreign DNA is always a factor.

Concatenated sequences of mitochondrial proteins coding genes (often translated) have been used extensively in animal molecular phylogenetics and have been shown to be especially useful for inferring demosponge relationships because of the low rate of mt-sequence evolution and a relatively homogeneous mtDNA nucleotide sequence composition in this group. Here we put together and analyzed a large dataset of mt-genomic sequences of demosponges, including 66 mtDNA sequences determined or assembled for this study. Importantly, we sampled all but one (Trachycladida) newly proposed orders within Heteroscleromorpha *sensu stricto*, the largest group of demosponges. We used this dataset to test the phylogenetic relationships and reconstruct the timing of major splits within the group.

### 4.2. Defining major lineages within Heteroscleromorpha

We are proposing four large groups (superorders) within Heteroscleromorpha *sensu stricto*:

- S1: freshwater haplosclerids (Spongillida), Scopalinida, and Sphaerocladina.
- S2: Axinellida, Biemnida, Bubarida along with *Topsentia ophiraphidites* and *Petromica* sp.
- S3: Tetractinellida: Astrophorina, Spirophorina, Rhizomorina (Sclero-todermidae, Azoricidae) + *Myceliospongia*.
- S4: Agelasida, Clionaida, Poecilosclerida, Polymastiida, Suberitida, and Tethyida.

We inferred the following relationships among these superorders: (S1(S2(S3,S4))). Our results are largely consistent with but add resolution to the study of Morrow and Cá rdenas (Morrow and Cardenas, 2015). The grouping of fresh-water haplosclerids with heteroscleromorphs rather than marine haplosclerids has been inferred in several previous studies (Borchiellini et al., 2004; Lavrov et al., 2008; Morrow et al., 2012; Redmond et al., 2013) using mtDNA and rRNA data and has been recently supported by a comprehensive genomic study (Simion et al., 2017). Similarly, grouping of *Vetulina* with freshwater sponges has been previously suggested (Redmond et al., 2013) and has been investigated in more details by Schuster *et al*. (Schuster et al., 2018a). It should be noted that the *Vetulina* sp. mtDNA sequence reported in the later study appears to be a chimera from several sponge species and was not included in the present analysis.

However, there are a few differences between this and previous molecular studies:

First, in our analysis Tetractinellida formed a sister group to S4 (Agelasida, Polymastiida, Clionaida, *etc*.) rather than Biemnida (Morrow et al., 2013, 2012) or S2 (Axinellida, Biemnida, Bubarida) (Morrow and Cardenas, 2015).

Second, within the S4, Tethyida grouped with Suberitida rather than Clionaida & Poecilosclerida (Morrow and Cardenas, 2015).

Third, within S2, Bubarida was a sister group to Biemnida, rather than Axinellida (Morrow and Cardenas, 2015).

Fourth, *Topsentia ophiraphidites* and *Petromica* sp. (classified as Suberitida and *incertae sedis*, respectively in (Morrow and Cardenas, 2015)) grouped together and formed a sister group to Axinellida. It has been noted that *Topsentia* appears to be a polyphyletic genus, with some species related to Axinellida, while other to Suberitida (Morrow et al., 2013). Because there are no molecular data from the type species of *Topsentia*, the proper classification of this genus remains undetermined.

Finally, *Svenzea flava* grouped with Dictyonellida (Bubarida) rather than with Scopalinida, which contained the type species *Svenzea zeai*. It has been reported the skeletal arrangement in *S. flava* differed to those in other members of the genus and that *S. flava* lacked the characteristic granular cells and large embryos/larvae found in the type species. Thus in the future *S. flava* will need to be renamed and reassigned to Dictyonellidae (Bubarida).

Our analysis also provided support for some of the redefined orders within Heteroscleromorpha. In particular, the redefined order Poecilosclerida, from which five families where reassigned to other orders, is now well supported by both sequence and gene order data.

### 4.3. Timing molecular phylogeny

In addition to resolving phylogenetic relationships among major lineages (orders) of heteroscromorphs, we provide molecular clock estimates for the divergences among them. While several molecular clock studies using demo-sponge mt-genome data have been recently conducted (Ma and Yang, 2016; Schuster et al., 2018b) they used much smaller datasets of mitogenomic sequences. Paradoxically, Schuster et al. (2018) study was also based on a clearly erroneous phylogenetic tree of demosponge relationships with marine haplosclerids placed deep within the G4 clade (compare to Lavrov et al., 2008; Simion et al., 2017), which made most of their time estimates *a priori* misleading.

A shared difficulty in molecular timing analysis of sponges is the scarcity of available calibration points. While the Paleozoic record of sponges is well established and extends to the Lowermost Cambrian (529-541 Ma) (Chang et al., 2017), with several lineages (e.g., the Vauxiidae, Anthaspidellidae and Hazeliidae) convincingly shown to belong to the class Demospongiae (Finks and Rigby 2004; Pisera 2004, 2006; Botting et al. 2013), the taxonomicm affinities of Paleozoic sponges are often controversial. Nevertheless, recent finding of well-preserved crown-group demosponges (Botting et al., 2015) constrain the upper bound for the basal split in demosponges at 515 MY.

The Precambrian record of sponges is even more controversial. Although there have been numerous reports of Precambrian Porifera fossils, most of them are not substantiated (Antcliffe et al., 2014). Instead, the current argument for the Precambrian origin of demosponges is based primarily on the presence of fossil steroids (in particular 24-isopropylcholestanes and recently discovered 26-methylstigmastane) in the geological record before the end of the Marinoan glaciation (635 MY ago) (Love et al., 2009; Zumberge et al., 2018). The problem with both of these biomarkers, however, has been their very spotty distribution across modern demosponges, which at best can be interpreted as a result of multiple independent losses (Zumberge et al., 2018). Such rampant loss makes it extremely difficult to estimate the origin of this synthetic pathway on a phylogenetic tree, especially given the sparse sampling of non-demosponge taxa (Zumberge et al., 2018). Furthermore, a recent study has shown that the biosynthesis precursors of “sponge biomarkers” are found among Rhizaria, heterotrophic protists common in the ancient and modern oceans (Nettersheim et al., 2019). Thus for the present study we used the beginning of Cambrian (541 MY) as the lower bound for the common ancestor of demosponges.

We used the origin of freshwater sponges (the split between freshwater sponges and Vetulina sp.) as the second calibration point for our analysis. The upper limit for the origin of freshwater sponges was defined by the oldest reported freshwater sponge fossils (Schindler et al., 2008). The lower bound was defined by a somewhat generic observation that species diversification in lakes prior to the Devonian was limited by low nutrient loads and high sediment loads (Cohen, 2003), There exist substantial uncertainty with both of these estimates. First, the fossils we used to define the upper limit of the split predate most other known freshwater sponge fossils (mostly Jurassic and Cretaceous) by more than 100MY (Pronzato et al., 2017). It is not clear if the lack of fossil freshwater sponges between the end of the Palaeozoic and the Jurassic is an artefact or if the colonization of the freshwater environment by sponges has really taken place more than once. Furthermore, although we are discussing the origin of freshwater sponges, in the analysis we constrained the time for the split between the lineages leading to *Vetulina* vs. freshwater sponges, which has likely happened in marine environment and thus preceded the transition to freshwater.

Our third calibration point was the origin of the crown group Baikal sponges (defined as the split between *Baikalospongia intermedia profundalis* and *Lubomirskia baicalensis*) and placed between 30 and 6 MY. The lower bound for this estimate is based on Lake Baikal age (estimated between 25-30 MY), and lack of Lubomirskiidae spicules in early Tertiary sediments of the Tunkinskaya land basin (approximately Oligocene or 23-33 MYA) (Veynberg, 2009). By contrast, studies of a part the core BDP corresponding to 6,50 – 4,75 MYA uncovered well-formed spicules of several species belonging to all four genera of the Lubomirskiidae family (Veynberg, 2009).

Given the substantial uncertainties in the calibration points, and a large variance associated with molecular clock analysis, divergence times estimates need to be taken with a grain of salt. Nevertheless, it is illuminating to see that most proposed orders of demosponges have an ancient (Mid-Paleozoic) origin. By contrast, the origin of crown groups within each of the order roughly corresponds to the Late-Paleozoic, or Mesozoic. We propose to use these time estimates to re-adjust the scopes of several orders within the Heteroscleromorpha *sensu stricto*.

Defining higher taxa in general and those of demosponges in particular is a tricky affair. While many zoologists associate higher taxa with substantial morphological disparity, measuring such disparity in sponges is difficult because of the paucity of morphological characters and high degree of convergent evolution. Using inferred times for various clades might provide a more objective way to define the ranks in this group. This also corresponds to the idea that true phenotypic divergence between lineages is a function of time. From this perspective, Merliida and Desmacellida should be united as a single order or even placed within the order Poecilosclerida. By contrast, the two Axinellida lineages represented in our study by *Axinella* and *Heteroxya* vs. *Stelligera* + *Ptilocaulis, Ectyoplasia*, and *Raspaciona* may need to be split into two orders (e.g., Axinenellida and Stelligerida as in (Morrow et al., 2012)). Finally, we recommend creating two new orders, one for *Topsentia ophiraphidites* and *Petromica* sp., the other for *Myceliospongia araneosa, Microscleroderma* sp., and *Leiodermatium* sp.

### 4.4. Phylogenetic position of Myceliospongia araneosa

*Myceliospongia araneosa*, the only described species in the genus *Myceliospongia* Vacelet & Perez, 1998, is one of the most unusual demosponges in its anatomy and cytology. The sponge is known from a single cave (the “3PP” cave near La Ciotat (43°09.47’N - 05°36.01’E)) in the Mediterranean, where it grows on vertical or overhanging in a cool stable environment. It has an encrusting ‘body’ of up to 25 cm diameter and 1 mm thick surrounded by a reticulation of filaments 5–x *μm* in diameter. These filaments form an extensive network around the body that can grow over and under other marine life. The sponge body has a reduced aquiferous system and a low number of small choanocyte chambers. Neither spicules nor spongin fibres are present, and collagen fibrils do not form bundles or thick condensations. The sponge also shows a remarkable uniformity of cell types, with the mesohyl cells being poorly differentiated. Because of the highly unusual organization, which offered little clues as to its affinities to other sponges, *Myceliospongia* was placed in Demospongiae, *incertae sedis* (Vacelet et al., 2002).

Phylogenetic analyses based both on sequence and gene order data have unequivocally placed *Myceliospongia* wihtin Tetractinellida, and as the sister group of Rhizomorina. The split between Myceliospongia + Rhizomorina and Astrophorina + Spirophorina has been estimated to occur 456 – 385 MYA, justifying establishing a new orders for these groups (as also suggested in (Schuster et al., 2017)).

### 4.5. Limitations of mitochondrial phylogenomics

While mt-genomic dataset has been predicted and proven informative for studying demosponge phylogenetic relationships, it is important to remember that mtDNA is inherited as a single locus (*e.g*., Avise, 2004, p. 74) so its history may be different from the species tree (Maddison, 1997). There are two main potential sources of gene tree vs. species tree incongruence for mtDNA data: incomplete lineage sorting (or deep coalescence) and mitochondrial introgression.

Incomplete lineage sorting refers to the retention of ancestral allelic polymorphism through a speciation event. If alternative alleles are than fixed in the two daughter species, the divergence time of the locus will precede the origin of the species. If allelic polymorphism is retained through two consecutive speciation events, the gene topology may become incongruent with the species topology. Because mitochondrial sequences tend to coalesce more quickly than nuclear ones (Moore, 1995) and because studies on extant species of demosponges demonstrated a remarkably low level of polymorphism in their mitochondrial genes (Duran et al., 2004; Wörheide, 2006; Ereskovsky et al., 2011), incomplete lineage sorting is not expected to have large influence on demosponge phylogenies.

Mitochondrial introgression refers to the horizontal acquisition of mtDNA via interspecific hybridization. While interspecific hybridization results in the transfer of nuclear as well as mitochondrial genes, mtDNA has been implicated in the vast majority of the reported cases of introgression in animals. Explanations for tendency of mtDNA to cross species boundaries include pure demography/drift, sex-biased interspecific mating, as well as adaptation (reviewed by Toews and Brelsford, 2012). Mitochondrial introgression and its influence on mtDNA-based phylogenies and identification have not been studied in demosponges. Theoretical consideration paint a contradictory picture. On the one hand, low level of sequence divergence among closely related species should simplify mitochondrial introgression by minimizing potential nuclear-mitochondrial incompatibilities in hybridization events. On the other hand, the same low level of sequence divergence will minimize possibility for positive selection and also render potential mitochondrial introgression little phylogenetically informative.

### 4.6. Future directions

The present study supported and expanded the study by Morrow et al. (Morrow and Cardenas, 2015) that proposed the first classification scheme for the class Demospongiae based on molecular data. A way forward in building a comprehensive classification for the Demospongiae is to extend the present sampling of taxa first to the family and, eventually, to the genus and species levels. We can expect that different phylogenetic markers will be more informative at different levels of demosponge phylogeny and that smaller datasets may be sufficient for resolving more shallow demosponge relationships (*e.g*., Gazave et al., 2013; Willenz et al., 2016). It is important, however, to use several independent molecular datasets for phylogenetic inferences, as this practice will help to recognize the artifacts of phylogenetic reconstruction and also to resolve the position of taxa that are problematic for each individual marker (*e.g*., Homoscleromorpha for nuclear rRNA and Dictyoceratida for mtDNA). It is also critical that the new molecular sampling is accompanied by comprehensive morphological studies on the vouchers. Finally, we want to emphasize that the resolved phylogeny is only the first step in understanding demosponge evolution as it provides the necessary framework for future studies of morphological, physiological, biochemical, and life history features of organisms as well as their geographical distribution.

## Acknowledgements

This work was supported by the National Science Foundation’s Assembling the Tree of Life program (DEB No. 0829783 to DVL, as well as DEB awards 0829763, 0829791, and 0829986), grant BFU2008-00227/BMC of the Spanish Ministry of Sciences and Innovation to MM, and by internal funds from Iowa State University. In addition, we gratefully acknowledge postdoctoral funding from the Irish Research Council to CCM. We thank our colleagues Steven Cook, Alexander Ereskovsky, April and Malcolm Hill, Gisele Lôbo-Hajdu, Jenna Moore, Shirley Pomponi, John Reed, Julie Reveillaud, Antonio Solé-Cava, Robert Thacker, and Gert Wörheide for samples contributed to this project and Cristina Diaz, Andrzej Pisera, and Sven Zea for help with species identification. We also thank the following graduate and undergraduate students in the Lavrov Lab for their help with the molecular work: Andrea Bekic, Jenessa Filler, Philip Lange, Katrina Lutap, Benjamin Sheller, Xiujuan Wang, and Katherine Wilson.

